# Decoding of emotion expression in the face, body and voice reveals sensory modality specific representations

**DOI:** 10.1101/869578

**Authors:** Maarten Vaessen, Kiki Van der Heijden, Beatrice de Gelder

## Abstract

A central issue in affective science is whether the brain represents the emotional expressions of faces, bodies and voices as abstract categories in which auditory and visual information converge in higher order conceptual and amodal representations. This study explores an alternative theory based on the hypothesis that under naturalistic conditions where affective signals are acted upon that rather than reflected upon, major emotion signals (face, body, voice) have sensory specific brain representations. During fMRI recordings, participants were presented naturalistic dynamic stimuli of emotions expressed in videos of either the face or the whole body, or voice fragments. To focus on automatic emotion processing and bypass explicit emotion cognition relying on conceptual processes, participants performed an unrelated target detection task presented in a different modality than the stimulus. Using multivariate analysis to asses neural activity patterns in response to emotion expressions in the different stimuli types, we show a distributed brain organization of affective signals in which distinct emotion signals are closely tied to the sensory origin. Our findings are consistent with the notion that under ecological conditions the various sensory emotion expressions have different functional roles, even when from an abstract conceptual vantage point they all exemplify the same emotion category.

## INTRODUCTION

In the course of daily social interactions, emotion signals from the face, voice and body are recognized effortlessly and responded to spontaneously when rapid adaptive actions are required. The specifics of the subjective experience in the natural environment determine which affective signal dominates and triggers the adaptive behavior. Rarely are the face, the whole body and the voice equally salient. That is, the actual conditions under which we react to an angry face may be different from those of hearing an angry voice or viewing whole body movements. For example, we see faces from close by and therefore personal familiarity may play a role in how we react to the angry face. This is less so for the voice or the whole body, both of which already prompt reactions when seen or heard from a distance while information about personal identity is not yet available or needed for action preparation. Thus, the angry body expression viewed from a distance and the angry face expression seen from closeby may each trigger a different reaction as behavior needs to be adapted to the concrete context. Therefore, a representation of affective meaning that is sensitive to the spatiotemporal parameters may seem desirable rather than an abstract system of higher order concepts as traditionally envisaged by emotion theorists (Ekman P and D Cordaro 2011); but see (Lindquist KA et al. 2012).

Research on the brain correlates of emotion has favored the traditional notion of abstract neural representations of basic emotions and this has also been the dominant rationale for multisensory research. Studies comparing how not just the face but also the voice and the whole body convey emotions have followed this overall basic emotion perspective and asked where in the brain affective information from different sensory systems converges (Peelen MV et al. 2010; Klasen M et al. 2011). Such representations were found in high-level brain areas known for their role in categorization of mental states (Peelen MV *et al.* 2010). Specifically, medial prefrontal cortex (MPFC) and superior temporal cortex (STS) represent emotions perceived in the face, voice or body at a modality-independent level. Furthermore, these supramodal or abstract emotion representations presumably play an important role in multisensory integration by driving and sustaining convergence of the sensory inputs towards the amodal emotion representation (Gerdes AB et al. 2014).

The present study explores a different perspective, complementary or compatible with that of high level abstract emotion representations yet motivated by the natural variability of daily emotion perception conditions that do not routinely involve conscious use of verbal labels. Emotion perception in naturalistic conditions is often driven by a specific context relative to a behavioral goal, as opposed to the conditions in the lab, where tasks often use explicit evaluation of emotional expressions. Under natural conditions, each sensory modality may have its own functionality such that e.g. fear is more effectively conveyed by the face, anger by the body and happiness by the voice. If so, brain responses would be characterized by specific emotion-modality combinations.

Additionally, the notion that supramodal representations of basic emotions are the pillars of emotion processing in the brain and allow for smooth convergence between the different sensory modalities is not fully supported by the literature. First, since the original proposal by Ekman (Ekman P 1992) and the constructivist alternative argued by Russell (Russell JA 2003) and most recently Feldman Barrett (Lewis M et al. 2010; Barrett LF 2017), the notion of a set of basic emotions with discrete brain correlates continues to generate controversy (Kragel PA and KS LaBar 2016; Saarimaki H et al. 2016). Second, detailed meta-analyses of crossmodal and multisensory studies, whether they are reviewing the findings about each separate modality or the results of crossmodal studies (Dricu M and S Fruhholz 2016; Schirmer A and R Adolphs 2017), provide a mixed picture. Furthermore, these meta-analyses also show that a number of methodological obstacles stand in the way of valid comparisons across studies. That is, taking into account the role of task (incidental perception, passive perception, and explicit evaluation of emotional expression) and the use of appropriate control stimuli reduces the number of studies that can validly be compared. Third, findings from studies that pay attention to individual differences and to clinical aspects reveal individual differences in sensory salience and dominance in clinical populations, for example in autism and schizophrenia. For example, (Karle KN et al. 2018) report an alteration in the balance of cerebral voice and face processing systems in the form of an attenuated face-vs-voice bias in emotionally competent individuals. This is reflected in cortical activity differences as well as in higher voice-sensitivity in the left amygdala. Finally, even granting the existence of abstract supramodal representations – probably in higher cognitive brain regions - it is unclear how they relate to early stages of affective processing where the voice, the face and the body information are processed by different sensory systems comprising distinct cortical and subcortical structures.

Here we used naturalistic dynamic stimuli to investigate whether the brain represents different sensory emotion expressions as modality specific or modality-invariant, using fMRI with dynamic stimuli expressing affect with either the body, the face or the voice. Importantly, we investigate these processes independently of the explicit evaluation of emotion to study whether the brain still differentiates between emotional expressions even when they are not in the direct focus of attention. We perform multivariate pattern analysis to identify cortical regions containing representations of emotion independently of the explicit evaluation of emotion.

For the sake of clarity, we contrast the implications of the centralist view and the distributed modality view for the neural representation of emotions. Following the first, there should be localized representations (here multi voxel patterns) for specific emotions that are independent of modality. These region(s) would show the following behavior: (1) respond to all stimuli above some threshold (i.e. be activated by all emotions/modalities); (2) have no discernable activation pattern between modalities, that is, modality cannot be decoded from the voxel activation patterns; (3) exhibit distinct voxel activation patterns for each emotion (emotion is decodable from the regions). This would provide strong evidence for a cross-modal or modality independent emotion representation and correspond to the classical notion of basic emotions in the literature. The alternative, distributed theory postulates that emotions are represented in the brain in a modality-emotion specific way. For example, some specific emotion-modality combination like a fearful body will elicit activation patterns that are different from fear from the voice or the face. Likewise, hearing a happy voice (laughing) can elicit different brain responses from seeing a happy expression from the face or body. In terms of testing this hypothesis, we would expect to find brain regions where emotion can be decoded but only within a specific modality, not across modalities. That pattern would provide clear evidence that the decoding will be driven by specific responses to emotion-modality combination.

## METHODS

### Participants

Thirteen healthy participants (mean age = 25.3; age range = 21-30; two males) took part in the study. Participants reported no neurological or hearing disorders. Ethical approval was provided by the Ethical Committee of the Faculty of Psychology and Neuroscience at Maastricht University. Written consent was obtained from all participants. The experiment was carried out in accordance with the Declaration of Helsinki. Participants either received credit points or were reimbursed with monetary reward after their participation in the scan session.

### Stimuli

Stimuli consisted of color video and audio clips of four male actors expressing three different emotional reactions to specific events (e.g. fear in a car accident or happiness at a party). Images were shown to the actors during recordings with the goal of triggering spontaneous and natural reactions of anger, fear, happiness and an additional neutral reaction. A full description of the recording procedure, the validation and the video selection is given in (Kret ME, S Pichon, J Grezes, et al. 2011). In total there were 16 video clips of facial expressions, 16 video clips of body expressions, and 32 audio clips of vocal expressions, half of which were recorded in combination with the facial expressions and half of which were recorded in combination with the body expressions (i.e. two audio clips per emotional expression per actor). All actors were dressed in black and filmed against a green background under controlled lighting conditions.

Video clips were computer-edited using Ulead, After Effects, and Lightworks (EditShare). For the body stimuli, faces of actors were blurred with a Gaussian mask such that only the information of the body was available. The validity of the emotional expressions in the video clips was measured with a separate emotion recognition experiment (emotion recognition accuracy > 80%). For more information regarding the recording and validation of these stimuli, see (Kret ME, S Pichon, J Grezes, *et al.* 2011; Kret ME, S Pichon, J Grèzes, et al. 2011).

### Experimental design and behavioral task

In a slow-event related design, participants viewed series of 1 second video clips on a projector screen or listened to series of 1 second audio clips through MR-compatible ear buds (Sensimetrics S14) equipped with sound attenuating circumaural earbuds (attenuation > 29 dB). The experiment consisted of 12 runs divided over 2 scan sessions. Six runs consisted of blocks of face and voice stimuli, followed by six runs consisting of body and voice stimuli. Blocks were either *auditory* (consisting of 18 audio clips) or *visual* (consisting of 18 video clips). These 18 trials within each block comprised 16 regular trials, and two catch trials requiring a response. Catch trials were included to determine that attention was diverted from explicit recognition or evaluation of the emotional expression by focusing attention on the other modality. That is, during visual blocks, participants were instructed to detect auditory catch trials, and during auditory blocks, participants were instructed to detect visual catch trials. For the auditory catch trial task, a frequency modulated tone was presented and participants had to respond whether the direction of frequency modulation was up or down. For the visual distractor task, participants indicated whether the fixation cross turned lighter or darker during the trial. A separate localizer session was also performed where participants passively viewed stimuli of faces, bodies, houses, tools and words in blocks; see (Zhan M et al. 2018) for details.

### Data acquisition

We measured blood-oxygen level-dependent (BOLD) signals with a 3 Tesla Siemens Trio whole body MRI scanner at the Scannexus MRI scanning facilities at Maastricht University (Scannexus, Maastricht). Functional images of the whole brain were obtained using T2*-weighted 2D echo-planar imaging (EPI) sequences [number of slices per volume = 50, 2 mm inplane isotropic resolution, repetition time (TR) = 3000 ms, echo time (TE) = 30 ms, flip angle (FA) = 90°, field of view (FoV) = 800 × 800 mm^2^, matrix size = 100 × 100, multi-band acceleration factor = 2, number of volumes per run = 160, total scan time per run = 8 min]. A three-dimensional (3D) T1-weighted (MPRAGE) imaging sequence was used to acquire high resolution structural images for each of the participants [1-mm isotropic resolution, TR = 2250 ms, TE = 2.21 ms, FA = 9°, matrix size = 256 × 256, total scan time = 7 min approx.]. The functional localizer scan also used a T2*-weighted 2D EPI sequence [number of slices per volume = 64, 2 mm in-plane isotropic resolution, TR = 2000 ms, TE = 30 ms, FA = 77, FoV = 800 × 800 mm^2^, matrix size = 100 × 100, multi-band acceleration factor = 2, number of volumes per run = 432, total scan time per run = 14 min approx.].

### Analysis

#### Pre-processing

Data were preprocessed and analyzed with BrainVoyager QX (Brain Innovation, Maastricht, Netherlands) and custom Matlab code (Mathworks, USA) (Hausfeld L et al. 2012; Hausfeld L et al. 2014). Preprocessing of functional data consisted of 3D motion correction (trilinear/sync interpolation using the first volume of the first run as reference), temporal high pass filtering (thresholded at five cycles per run), and slice time correction. We co-registered functional images to the anatomical T1-weighted image obtained during the first scan session and transformed anatomical and functional data to the default Talairach template.

#### Univariate analysis

We estimated a random-effects General Linear Model (RFX GLM) with a predictor for each stimulus condition of interest (12 conditions in total): four emotion conditions times three modalities (face, body, voice). We note that modality is used here in a broader sense than just the physical nature of the stimuli (sound versus visual). Additionally, we included predictors for the trials indicating the start of a new block and the catch trials. Predictors were created by convolving stimulus predictors with the canonical hemodynamic function. Finally, we included six motion parameters resulting from the motion correction as predictors of no interest. For this analysis, data was spatially smoothed with a 6mm full-width half-maximum (FWMH) Gaussian kernel. To assess where in the brain the two different experimental factors had an influence, an ANOVA was run with either modality or emotion as a factor. Additionally the ANOVA with the emotion factor was run for each modality separately. As effect sizes were generally low, the final group statistical maps were liberally thresholded at p<0.001 uncorrected. For visualization purposes, the group volume maps were mapped to the cortical surface. As this operation involves resampling the data (during which the original statistical values get lost), surface maps are displayed with discrete label values instead of continuous statistical values. As such, we do not include a colorbar in the surface map figures.

#### Multivariate analysis

We first estimated beta parameters for each stimulus trial with custom MATLAB code by fitting an HRF function with a GLM to each trial in the time series. These beta values were then used as input for a searchlight multivariate pattern analysis (MVPA) with a Gaussian Naïve Bayes classifier (Ontivero-Ortega M et al. 2017). The searchlight was a sphere with a radius of 5 voxels. The Gaussian Naïve Bayes classifier is an inherently multi-class probabilistic classifier that performs similar to the much-used Support Vector Machine classifier in most scenarios, but is computationally more efficient. The classifier was trained to decode (1) stimulus modality (visual or auditory) (2) stimulus type (i.e. body, face or voice); (3) stimulus emotion (e.g. fear in all stimulus types vs angry in all stimulus types); (4) within-modality emotion (e.g. body angry vs. body fear) (5) cross-modal emotion (e.g. classify emotion by training on body stimuli and testing on the voice stimuli from the body session). Classification accuracy was computed by averaging the decoding accuracy of all folds of a leave one-run out cross-validation procedure. We tested the significance of the observed decoding accuracies at the group level with a one-sample t-test against chance (50% for modality, 33% for stimulus types, 25% for emotion). We also tested the emotion effect with an additional analysis where the neutral condition was excluded. As in the univariate analyses, these maps were transformed to the cortical surface for display purposes. The cross-modal decoding revealed several subcortical structures that are not visualized well on the cortical surface and therefor are displayed on a volume map (p<0.01, uncorrected).

To evaluate the relative contribution of either stimulus type or of emotion in terms of information content that can be decoded, we performed an analysis where all stimuli for a specific combination of two emotions (e.g. anger and happy from the same session) were extracted and a decoder was trained to either classify the two different emotions or the two different stimulus types (body/voice or face/voice). The resulting accuracy maps were first thresholded such that regions where both the stimulus type and emotion decoder accuracy were below chance at the group level were excluded. Next we contrasted the two accuracy maps at the group level i.e. accuracy for emotion > accuracy for stimulus type and displayed these as a volume map at p<0.001 uncorrected.

#### ROI analysis

Lastly, to gain more insight into details of the responses in some of the regions identified by the previously mentioned analyses, as well as regions known to be important for emotion or multi-modal integration (Peelen MV *et al.* 2010), we extracted beta values from these ROIs and made several plots that (1) display the decoding accuracy of the selected voxels for stimulus type and emotion, (2) display the mean beta values for each of the 12 conditions, (3) display the multivoxel representational dissimilarity matrix (Kriegeskorte N et al. 2008) and (4) warp the multivoxel patterns to a 2-dimensional space with multidimensional scaling and display the relative distance between the conditions with graphical indicators for the stimulus type (icon) and emotion (line color)

## RESULTS

### Univariate analysis

We ran an ANOVA with factors ‘stimulus type’ and ‘emotion’ on the beta values estimated with an RFX GLM on the entire data set, as well as separate ANOVAs with factor ‘emotion’ on the data of each stimulus type (i.e. faces, bodies, and voices). As expected, the F-map for stimulus type (see Fig. S1) revealed significant clusters with differential mean activation across stimulus types in primary and higher-order auditory and visual regions, as well as in motor, pre-motor and dorsal/superior parietal cortex. The univariate analyses did not reveal cortical regions showing a significant effect for emotion. Although the F-maps of these emotion analyses showed some small clusters at lenient thresholds (*p* < 0.001, uncorrected), none of these survived correction for multiple comparisons. See Supplementary results for details.

### Multivariate analysis with the GNB decoder

We aimed to decode stimulus type by training the classifier separately for the two sessions to decode visual vs. auditory stimuli: body vs. voice (session 1) and face vs. voice (session2). As expected, stimulus type could be decoded significantly above chance level (50%, p<0.05 FWE corrected) in auditory, visual and fusiform cortex, and large part of the lateral occipital and temporo-occipital cortex, presumably including the extrastriatal body area (EBA; see Fig. 1 top panel).

**Figure 1:**
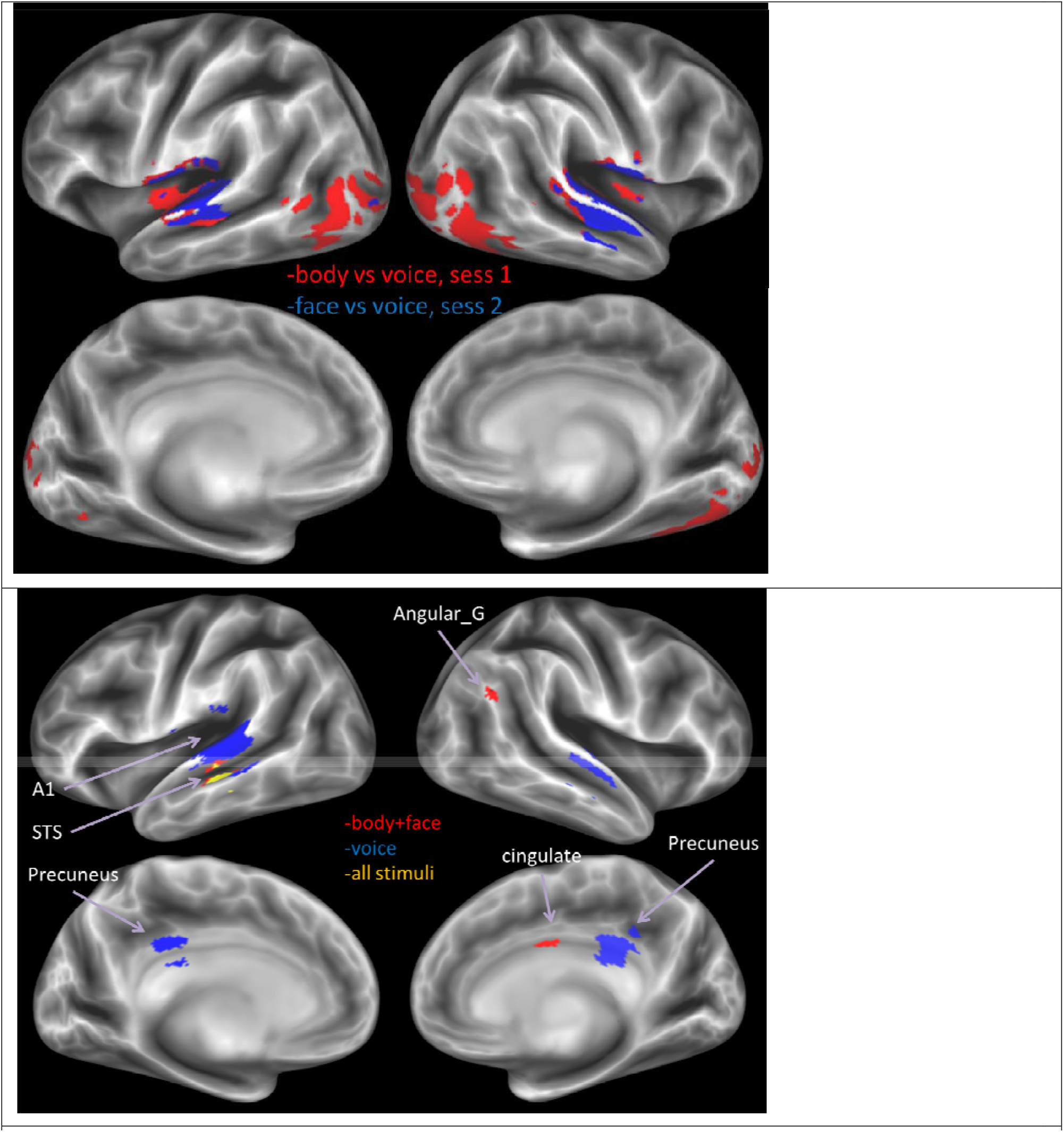
(Top) Decoder trained to classify stimulus type, for the two sessions separately, results derived from thresholding the volume map at p<0.05, FWE corrected. (Bottom) Decoder trained to classify emotion from all stimuli (yellow), all visual stimuli (face+body, red) and all voice stimuli (blue), The displayed results are label maps derived from the volume map thresholded at p<0.001 uncorrected with a minimum cluster size threshold of 3k=25 voxels. Abbreviations: Angular G = angular gyrus, A1 = primary auditory cortex, STS = superior temporal cortex.

Decoding of emotion resulted in qualitatively lower accuracies and smaller clusters compared to the decoding of modality. Above chance accuracies (33%) for decoding emotion from all stimulus types together were observed in STS (Fig. 1, bottom panel). Next, we trained and tested the classifier to decode emotions within a specific stimulus modality, that is, considering either the combined face and body stimuli (i.e. visual), or the voice stimuli (i.e. auditory). For body and face this revealed STS, cingulate gyrus and angular gyrus. Emotion could be decoded for voice stimuli in primary and secondary auditory regions (including the superior temporal gyrus [STG]), and the precuneus (see Fig 1 bottom panel). See Table 1 for details of these results.

**Table 1:** Results for the decoding of emotion.

|  | *cluster*<br>*size* | *clusgter*<br>*p(unc)* | *peak T* | *peak*<br>*p(unc)* | *x* | *y* | *z* |
| --- | --- | --- | --- | --- | --- | --- | --- |
| **All stimuli** |  |  |  |  |  |  |  |
| STS | 38 | 0.0536 | 5.8422 | 0.0000 | -3 | 21 | -7 |
| **Voice stimuli only** |  |  |  |  |  |  |  |
| planum temporale | 749 | 0.001 | 6.462 | 0.000 | -58 | -34 | 6 |
| posterior cingulate gyrus / precuneus | 187 | 0.058 | 5.654 | 0.000 | 8 | -34 | 30 |
| planum temporale | 210 | 0.046 | 5.206 | 0.000 | 62 | -18 | 2 |
| **Face and body stimuli** |  |  |  |  |  |  |  |
| STS | 59 | 0.0200 | 6.8069 | 0.0000 | -52 | -26 | -12 |
| right caudate | 92 | 0.0052 | 6.5513 | 0.0000 | 16 | -12 | 20 |
| angular gyrus | 27 | 0.0979 | 6.4339 | 0.0000 | 60 | -50 | 34 |

We also trained and tested the classifier to decode emotions within each visual category, that is, considering either the face or the body. This did not reveal any clusters where emotion could be decoded accurately (suprathreshold at p<0.05 FWE corrected), although additional results were obtained at more lenient thresholds (see Fig. S4).

### ROI Analysis

In addition to the whole-brain searchlight analysis, we performed a more sensitive region-of-interest (ROI) analysis. Specifically, we used data of an independent localizer (see *Methods*) to identify early auditory cortex, early visual cortex, rEBA, and rFFA. We furthermore included two multisensory regions, pSTS and mPFC that were previously identified as regions holding supramodal representations of emotion (Peelen MV *et al.* 2010). We used an anatomical definition of these areas, defined by a spherical ROI with a radius of 5 voxels centered on the reported cluster peak locations. Finally, we included the amygdala given its important role in previous studies, most recently in the study by (Whitehead JC and JL Armony 2019) using the univariate contrast face fear > face neutral (p<0.01 uncorrected).

In all ROIs, stimulus type could be decoded above chance level (one sample t-test against chance level, all p<0.0001). Furthermore, when the classifier operated on all stimuli together (that is, face, body, and voice), emotion could be successfully decoded in the EBA, auditory cortex, and left amygdala (all p<0.02). That is, in line with the results of the searchlight analysis, when the classifier operated on the data of each modality in isolation, decoding accuracies for emotion were above chance level in early visual cortex for face stimuli (p<0.03), and in auditory cortex for voice stimuli (p<0.002). Emotion could not be decoded above chance level in the supramodal regions (mPFC and pSTS). Taken together, these results demonstrate that these regions could not be identified as supramodal as the responses were not invariant to stimulus type (see Fig. 2). Qualitatively, the RDM and MDS plots in Figure 2 show that in all tested ROIs there is a strong effect of stimulus type (blocks on diagonal for the RDM and a large distance between types in the MDS). Notable, this effect is not always clearly present in just the beta magnitudes (response levels) and is at least partially caused by the multi-voxel pattern dissimilarities.

**Figure 2:**
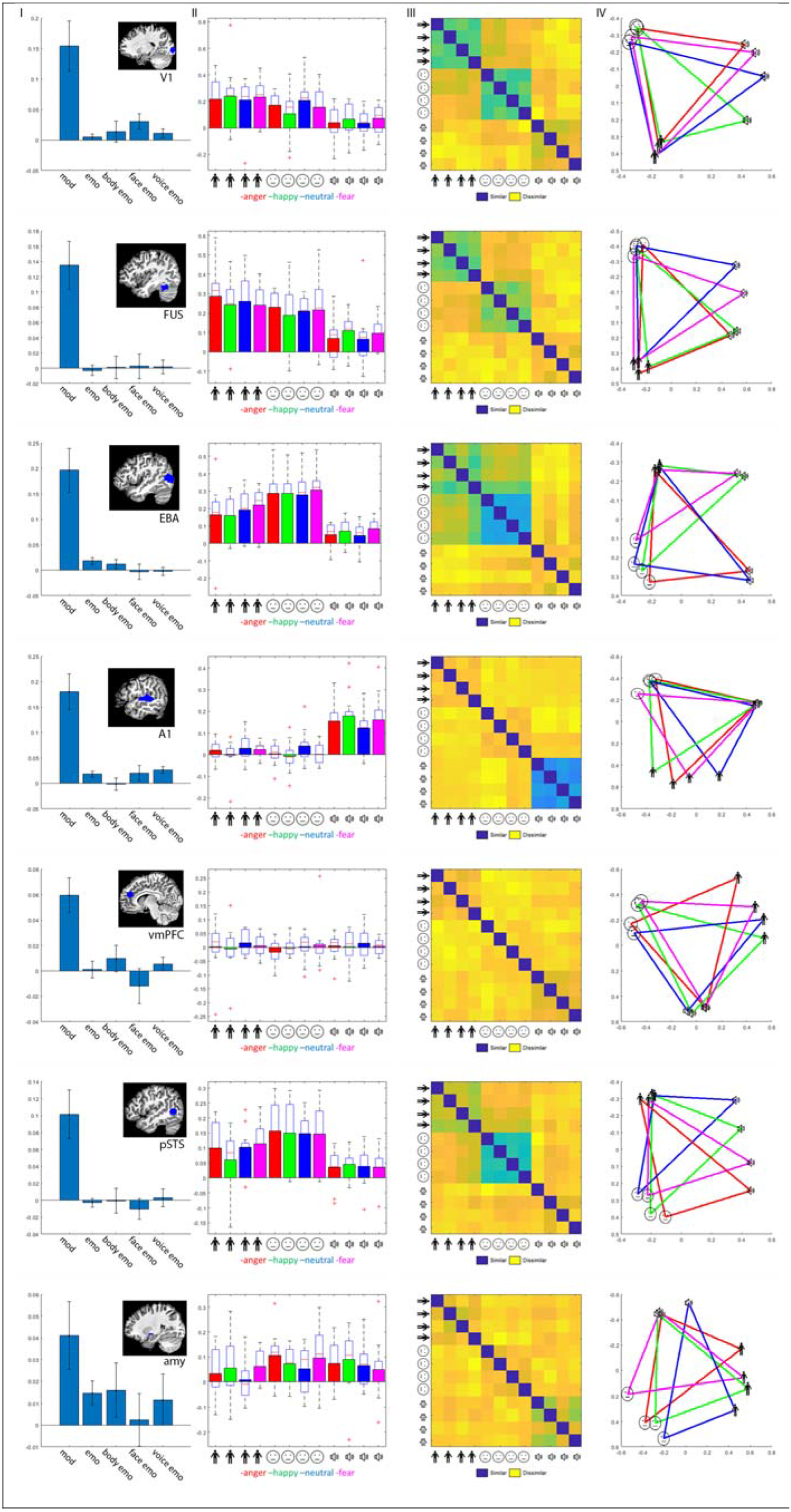
I) Location of the ROI and graphs indicating decoding accuracy for stimulus type and emotion and emotion for each stimulus type in the ROI. II) Trial-wise beta values for all 12 conditions (3x modality and 4x emotion) averaged over the ROI and group. Error bars indicate SE. II) Representational dissimilarity matrix for the ROI. IV) Multidimensional scaling plot of the group averaged trial-wise beta values. Line colors indicate same emotion, icons display modality. Distance in the plot is related to similarity of the ROI voxel activation patterns. Colors as in the beta plot for emotion.

### Cross-modal decoding

We performed an additional analysis to gain insight into where in the brain supramodal emotion regions might be found by training a classifier to decode emotion across stimulus modalities. Being able to predict emotion by training on one modality and testing on another modality would be a strong indication of supramodal emotion encoding in the brain. Therefore, the cross-modal classifier was trained (or tested) on either the body or face stimuli and tested (or trained) on the voice stimuli from the body or face session, respectively. Thus, four classifiers were trained in total (training on body and testing on voice, training on voice and testing body, training on face and testing on voice, training on voice and testing on face). In contrast to the successful decoding of modality and emotion within modality, none of these cross-modal classifiers resulted in accurate decoding of emotion (p<0.001, see Supplementary Fig. S5).

### Contrasting accuracies for emotion and modality

There was little overlap in regions identified by the within stimulus type classifier (see Fig. 2 and S4). Specifically, regions where emotion was decoded successfully from body stimuli did not converge with regions where emotion was decoded successfully from the face and/or voice. Therefore, to identify regions that have a purely supramodal representation of emotion, we contrasted the accuracy map for modality versus emotion directly. Here, finding regions having higher decoder accuracy for emotion compared to modality would be a strong indication for supramodal emotion encoding. This analysis revealed in general that, as expected, modality could be decoded with greater accuracy than emotion in primary auditory and visual regions, as well as in parietal and pre-motor regions. We could not clearly identify any regions where binary combination of two specific emotions could be decoded with higher accuracy than the modality. Only for fear and neutral in the face-voice session did we find a region in the medial motor cortex and for happy and neutral in the body-voice session in the white matter (at p<0.001 uncorrected, see Fig. S6).

## DISCUSSION

Our results indicate that the brain regions involved in emotion processing are modality specific. That is, in the regions where emotion can be decoded, it could only be decoded within modality but not across modalities, indicating that the decoding is driven by a specific response to a specific emotion-modality combination. In a departure from the few previous studies using a partially comparable approach we found evidence for sensory specific rather than abstract supramodal representations that sustain perception of various affective signals as a function of the modality (visual or auditory) and the stimulus category (face, voice or body). Because of the three stimulus categories used, the different emotions studied, the task conditions, and the converging results from different analysis techniques, our study presents a novel approach to understanding the specific contribution and the neural basis of emotional signals provided by different sensory modalities.

To understand our findings against the background of the literature, some specific aspects of our study must be highlighted. We used dynamic realistic face and body stimuli instead of point light displays or static images. The latter are also known to complicate comparisons with dynamic auditory stimuli (Campanella S and P Belin 2007). Next, our stimuli do not present prototypical emotion representations obtained by asking actors to portray emotions but present spontaneous whole body reactions to images of familiar events. The images we used may therefore be more spontaneous and trigger more sensorimotor processes in the viewer than posed expressions. Third, many previous studies used explicit emotion recognition (Lee KH and GJ Siegle 2012), passive viewing (Winston JS et al. 2003), implicit tasks like gender categorization (Dricu M and S Fruhholz 2016) or oddball tasks presented in the same modality as the stimulus. In contrast, our modality specific oddball task is presented in the alternate modality of the stimulus presentation thereby diverting attention not only from the emotion content but also from the perceptual modality in which the target stimuli of that block are shown. This task was intended to approximate the naturalistic experience of emotional signals, where often one is engaged in one activity (visual perception) when another event intrudes (an auditory event). We discuss separately the findings on the major research questions.

### Univariate analysis

Although our goal was to characterize neural responses with MVPA techniques, for the sake of comparisons with the literature, we also briefly discuss our univariate results. How do these results compare to findings and meta-analyses in the literature? As a matter of fact, there are no previous studies that used comparable materials (four emotion categories, three stimulus types, two modalities) and an other modality centered task like the present. The studies that did include bodies used only neutral actions, not whole body emotion expressions (Dricu M and S Fruhholz 2016) except for one study comparing face and body expression videos by (Kret ME, S Pichon, J Grezes, *et al.* 2011). Only the study by Peelen et al. used faces, bodies, and voices, but with a very different task as we discuss below (Peelen MV *et al.* 2010).

Compared to the literature, the findings of the univariate analysis present correspondences as well as differences. A previous study (Kret ME, S Pichon, J Grezes, *et al.* 2011) with face and body videos used only neutral, fear and anger expression and a visual oddball task. They reported that EBA and STS show increased activity to threatening body expressions and FG responds equally to emotional faces and bodies. For the latter, higher activity was found in cuneus, fusiform gyrus, EBA, tempo-parietal junction, superior parietal lobe, as well in as the thalamus while the amygdala was more active for facial than for bodily expressions, but independently of the facial emotion. Here we replicate that result for faces and bodies and found highly significant clusters with differential mean activation across stimulus types in primary and higher-order auditory and visual regions, as well as in motor, pre-motor and dorsal/superior parietal cortex (Fig. S1).

Regions sensitive to stimulus category were not only found in primary visual and auditory cortex as expected but also in motor, pre-motor and dorsal/superior parietal cortex consistent with the findings in Kret et al 2011. In view of the literature on perception of emotion expressions in either the face, voice or body it is not surprising that dorsal parietal cortex, pre- motor cortex and anterior insula (Fig. S2) differentially respond to emotions as different emotions trigger different adaptive actions (de Gelder B 2006; Grezes J et al. 2007; Pichon S et al. 2008; Whitehead JC and JL Armony 2019). Interestingly, the interaction effect between modality and emotion also revealed the retrosplenial cortex (Fig. S3). Retrosplenial cortex receives input from areas known to play a role in processing salient information (prefrontal cortex, superior temporal sulcus, precuneus, thalamus, and claustrum (Maddock RJ 1999). The retrosplenial cortex is associated with navigation and memory functions and may be part of a network that conveys predatory threat information to the cerebral cortex (de Lima MAX et al. 2019). A similar functionality may be reflected in this activity here. Activity in this area is also consistent with the recent findings that the retrosplenial cortex contributed to the accurate classification of fear stimuli. (Caruana F et al. 2018).

To summarize, this univariate analysis including three stimulus modalities and four emotion categories replicates some main findings about brain areas involved respectively in face, body and voice expressions while also revealing parietal and motor area activity but does not provide evidence for overlap in brain activity neither for modality nor emotion category.

### Multivariate analysis

The goal of our multivariate approach was to reveal the areas that contribute most strongly to an accurate distinction between the modalities and the stimulus emotion. Our results of the MVPA searchlight show that modality type can be decoded from the sensory cortex and that emotion can be decoded in STG for voice stimuli and in STS for face and body stimuli. We found no overlap in brain regions that contribute to the classification of emotion when using either the face, the body or the voice decoder (Fig. S4). Thus the brain areas that are involved in discriminating between face, voice or body expressions irrespective of the emotion are different from each other. Nor did we find evidence for neural activity overlap the other way around when using a cross-modal emotion decoder and looking for possible brain areas common to the modalities (Fig. S5). Lastly, we could not clearly identify regions where emotion could be decoded with higher accuracy than modality (Fig. S6). To put it negatively, we could not clearly identify supramodal emotion regions, defined by voxel patterns where emotion could be decoded and that would show very similar voxel patterns for the same emotion in the different modalities. This clearly indicated that the brain responds to facial, body and vocal emotion expression in a unique fashion. Thus the overall direction pointed to by our results seems to be that that being exposed to emotional stimuli (that are not task relevant and while performing a task requiring attention to the other modality than that in which the stimulus is presented) is associated with brain activity that shows both an emotion specific and a modality specific pattern.

### ROI analysis

To follow up on the whole-brain analysis we performed a detailed and specific analysis of a number of ROIs. For the ROIs based on the localizer scans (early visual, auditory areas as well as FFA and EBA) stimulus type could be decoded and these results are consistent with the MVPA searchlight analysis. We also defined ROIs based on the literature with the goal of comparing our results to findings about higher order areas that had been identified as supreamodal areas, mPFC and pSTS. This analysis revealed accurate decoding of stimulus type but no evidence of supramodal representations.

Two previous MVPA studies addressed similar issues investigated in this study using faces, bodies and voices (Peelen MV *et al.* 2010) or bodies and voices (Whitehead JC and JL Armony 2019). The first study reported medial prefrontal cortex (MPFC) and posterior superior temporal cortex as the two areas hosting supramodal emotion representations. These two areas do not emerge in our MVPA searchlight analysis. To understand this very different result it is important to remember that in their study participants were encouraged to actively evaluate the perceived emotional states. The motivation was that explicit judgments would increase activity in brain regions involved in social cognition and mental state attribution (Peelen MV *et al.* 2010). In contrast, the motivation of the present study was to approximate naturalistic perception conditions. Our design and task were intended to promote spontaneous non-focused processes of the target stimuli and did not promote amodal conceptual processing of the emotion content. It is likely that using an explicit recognition task would have activated higher level representations e.g. posterior STS, prefrontal cortex and posterior cingulate cortex that would then feedback to lower level representations and modulate these towards more abstract representations (Schirmer A and R Adolphs 2017). Note that no amygdala activity was reported in that study. The second study using passive listening or viewing of still bodies and comparing fear and neutral expressions also concludes about a distributed network of cortical and subcortical regions responsive to fear in the two stimulus types they used (Whitehead JC and JL Armony 2019). Of interest is their finding concerning the amygdalae and fear processing. While in their study this is found across stimulus type for body and voice, the classification accuracy when restricted to the amygdalae was not significantly above chance. They concluded that fear processing by the amygdalae heavily relies on contribution of a distributed network of cortical and subcortical structures.

Our findings suggest a different and a novel perspective on the role of the different sensory systems and the different stimulus categories that convey affective signals in daily life and fits with the role of emotions as seen from an evolutionary perspective. Our results are compatible with an ecological and context sensitive approach to brain organization (Cisek P and JF Kalaska 2010; Mobbs D et al. 2018) rather than with an exclusive focus on high order representation of emotion categories grounded in concepts and verbal labels. For comparison, a similar approach not to emotion concepts but to cognitive concepts was argued by Barsalou et al. (Barsalou LW et al. 2003). This type of distributed organization or emotion representation may be more akin to what is at stake in the daily experience of affective signals and how they are flexibly processed for the benefit of ongoing action and interaction in a broader perspective of emotions as states of action readiness (Frijda NH 2004).

Our results are relevant for two longstanding debates in the literature, one on the nature and existence of abstract emotion representations and basic categories and the other on processes of multisensory integration. Concerning the first one, our results have implications for the debate on the existence of basic emotions (Ekman P 2016). Interestingly, modality specificity has rarely been considered as part of the issue as the basic emotion debate largely focusses on facial expressions. The present results might be viewed as evidence in favor of the view that basic emotions traditionally understood as specific representations of a small number of emotions with an identifiable brain correlate (Ekman P 2016) simply do not exist but that these are cognitive-linguistic constructions (Russell JA 2003). On the one hand, our results are consistent with critiques of basic emotions theories and meta-analysis (Lindquist KA *et al.* 2012) as we find no evidence for representations of emotions in general or specific emotions within or across modality and stimuli. Affective information processing thus appears not organized as categorically, neither by conceptual emotion category nor by modality, as was long assumed. Emotion representation, more so even than object representation, may possibly be sensory specific or idiosyncratic (Peelen MV and PE Downing 2017) and neural representations may reflect the circumstances under which specific types of signals are most useful or relevant rather than abstract category membership. This pragmatic perspective is consistent with the notion that emotions are closely linked to action and stresses the need for more detailed ethological behavior investigations (de Gelder B 2016).

While our study was not addressing issues of multisensory perception, our findings may have implications for theories of multisensory integration. As has often been noted, human emotion research by and far is still limited to the study of facial expressions. In line with the dominant view studies extending the scope of facial expression based theories have primarily been motivated to discover similarities across different modalities and stimulus types. Our group has initiated studies that go beyond the facial expression and found rapid and automatic influence of one type of expression on another (face and voice, (de Gelder B et al. 1999); face and body (Meeren HK et al. 2005); face and scene (Righart R and B de Gelder 2008; Van den Stock J et al. 2013); body and scene, (Van den Stock J et al. 2014); auditory voice and tactile perception (de Borst AW and B de Gelder 2017). These original studies and subsequent ones (Müller VI et al. 2012) investigated the impact of one modality on the other and targeted the area(s) where different signals converge. For example, Müller et al. (Müller VI *et al.* 2012) report posterior STS as the site of convergence of auditory and visual input systems and by implication, as the site of multisensory integration. Note that here and in many other studies the amygdala and its connectivity to face and voice areas emerges as a core structure involved in multisensory integration. Current studies did not yet raise the question of sensory specificity of the representations at stake in such cross-modal effects. Our findings stress the importance of sensory specific representation and indicate that aside from loci of integration based presumably on amodal representations, multisensory perception and integration seems to respect stimulus complementarity in the vertical plane rather than convergence between different emotion signals onto an abstract supramodal representation.

The motivation to include three stimulus categories led to some limitations of the current design because two separate scanning sessions were required to have the desired number of stimulus repetitions. To avoid that the comparison of representations of stimuli from two different sessions was biased by a scan session effect, we did not include any results that referred to differences or commonalities of stimuli from different sessions (e.g. bodies vs faces). A second limitation is that the voice stimuli were not controlled for low level acoustic features over different emotions and therefore results from the decoding of emotion from the voice stimuli may partly have reflected these differences.

### Conclusions

Our results show that the brain correlates of observing emotional signals from the body, the face of the voice are specific for the modality as well as for the specific signal within the same modality. Our results underscore the importance of considering the specific contribution of each modality and each type of affective signal rather than only their higher order amodal similarity. We suggest that future research may look into the differences between the emotion signals and how they are complementary and not only at amodal similarity.

## Acknowledgments

We would like to thank Dr. Rebecca Watson for acquiring the MRI data. This work was supported by the European Research Council (ERC) FP7-IDEAS-ERC Grant agreement number 295673 *Emobodies*, by the Future and Emerging Technologies (FET) Proactive Programme H2020-EU.1.2.2 Grant agreement 824160 *EnTimeMent* and by the Industrial Leadership Programme H2020-EU.1.2.2 Grant agreement 825079 *MindSpaces*.

## Supplementary results

### Univariate

The F-map for emotion showed regions with different activation levels across emotions in insular cortex, the superior temporal sulcus (STS), retrosplenial cortex, angular gyrus, and superior medial occipital cortex, see Fig. S2.

We ran separate ANOVAs with factor ‘emotion’ for each stimulus type condition (voices, faces, bodies; see Fig. S2), to identify regions that respond differentially across emotions within a specific stimulus type. For the face condition, this revealed clusters in STS and retrosplenial cortex. For the body condition, we observed regions in inferior temporal cortex and the intraparietal sulcus (IPS). Finally, the ANOVA for the voice condition (for session 1 and session 2 separately) revealed auditory cortex, premotor cortex, IPS and angular gyrus.

We then tested for an interaction effect between the factors stimulus type and emotion. This revealed clusters in the retrosplenial cortex, the dorsolateral prefrontal cortex (dLPFC), auditory cortex and insula, see Fig. S3.

**Figure S1:**
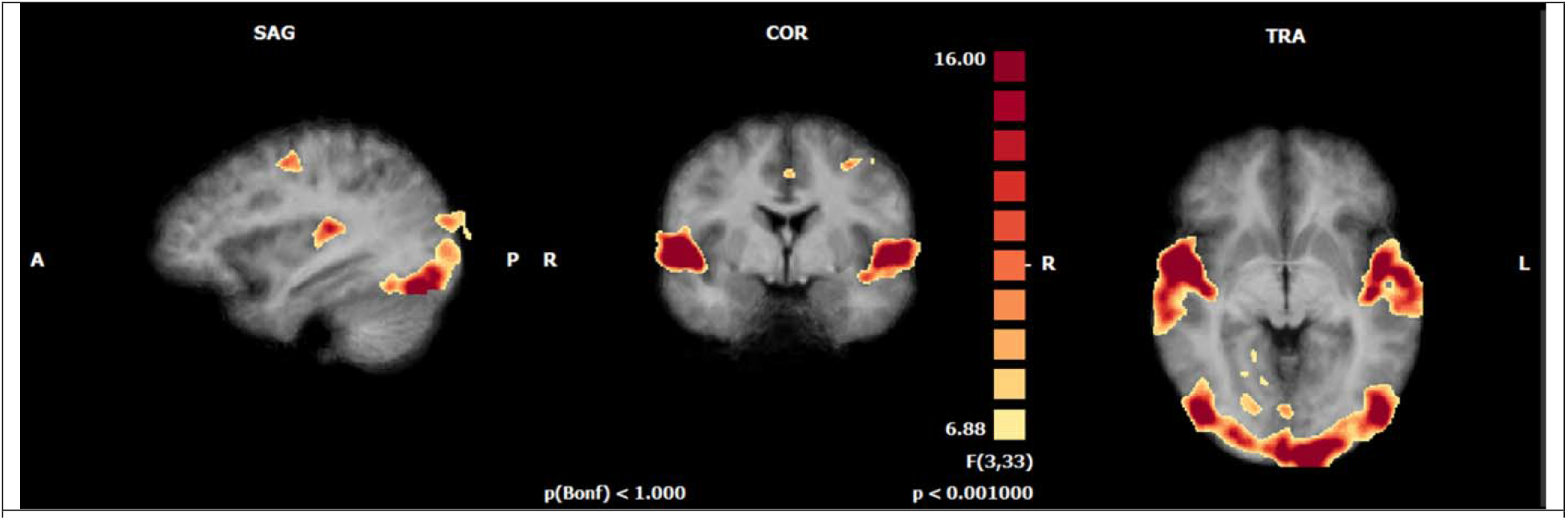
F-map (p<0.001 uncorrected) for stimulus type effect

**Figure S2.**
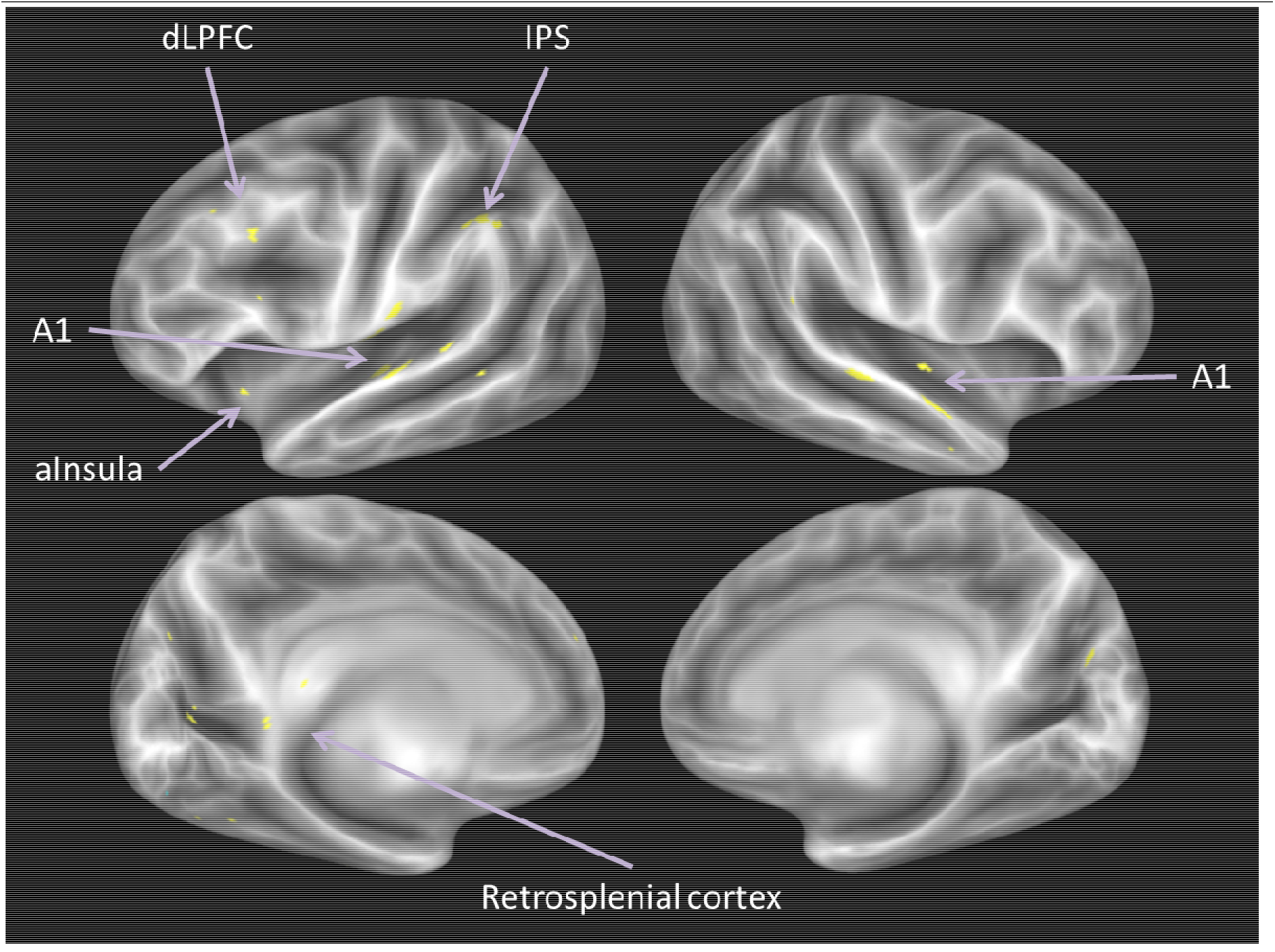
F-map for the ANOVA with factor ‘emotion’, calculated either for all stimuli (orange) or for each stimulus type separately (bodies = red; faces = green; voices session 1 = dark blue; voices session 2 = light blue). The displayed results are label maps derived from the volume map thresholded at p<0.001 uncorrected.

**Figure S3.**
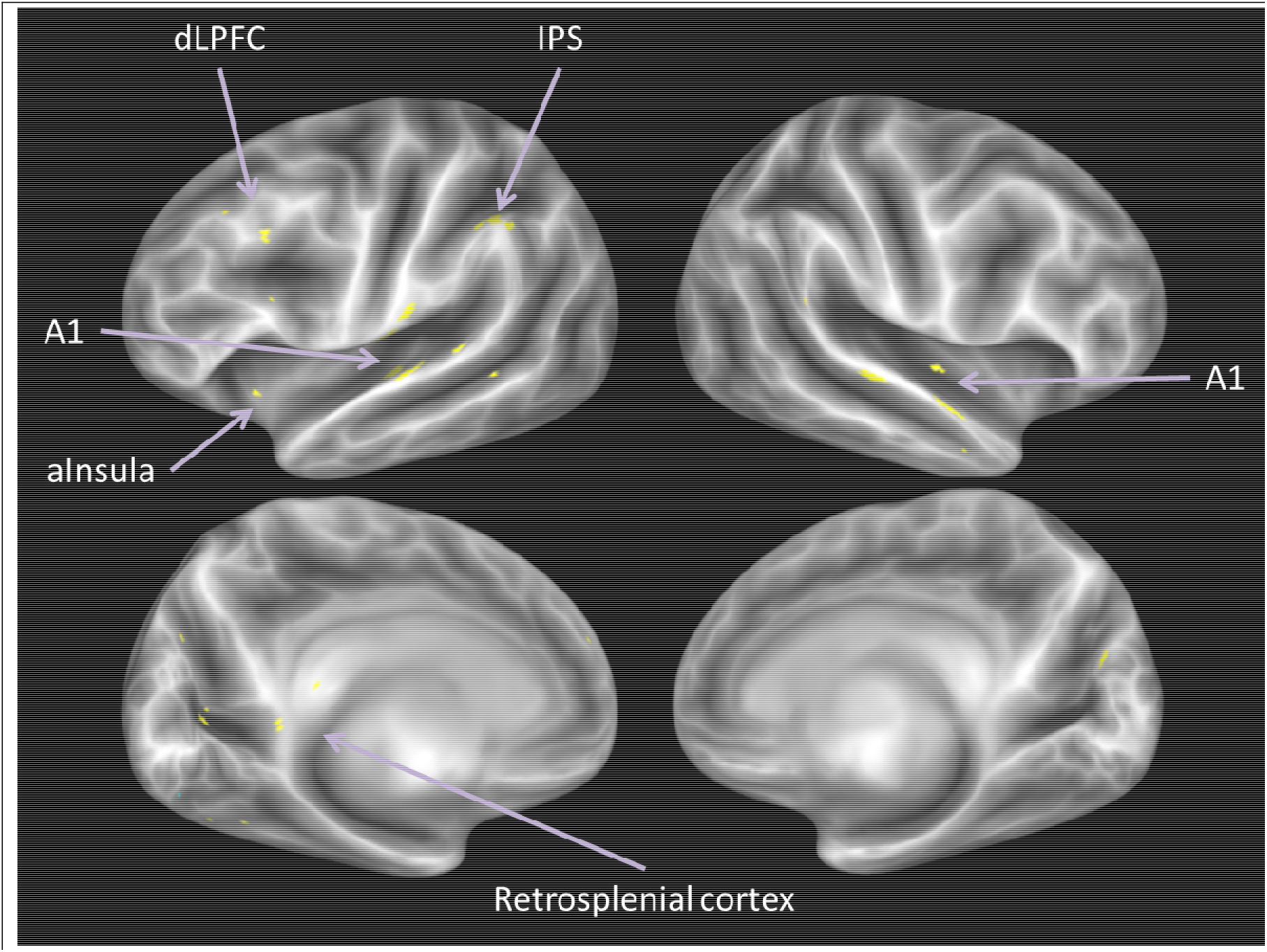
Interaction emotion and stimulus type. The displayed results are label maps derived from the volume map thresholded at p<0.001 uncorrected.

**Figure S4:**
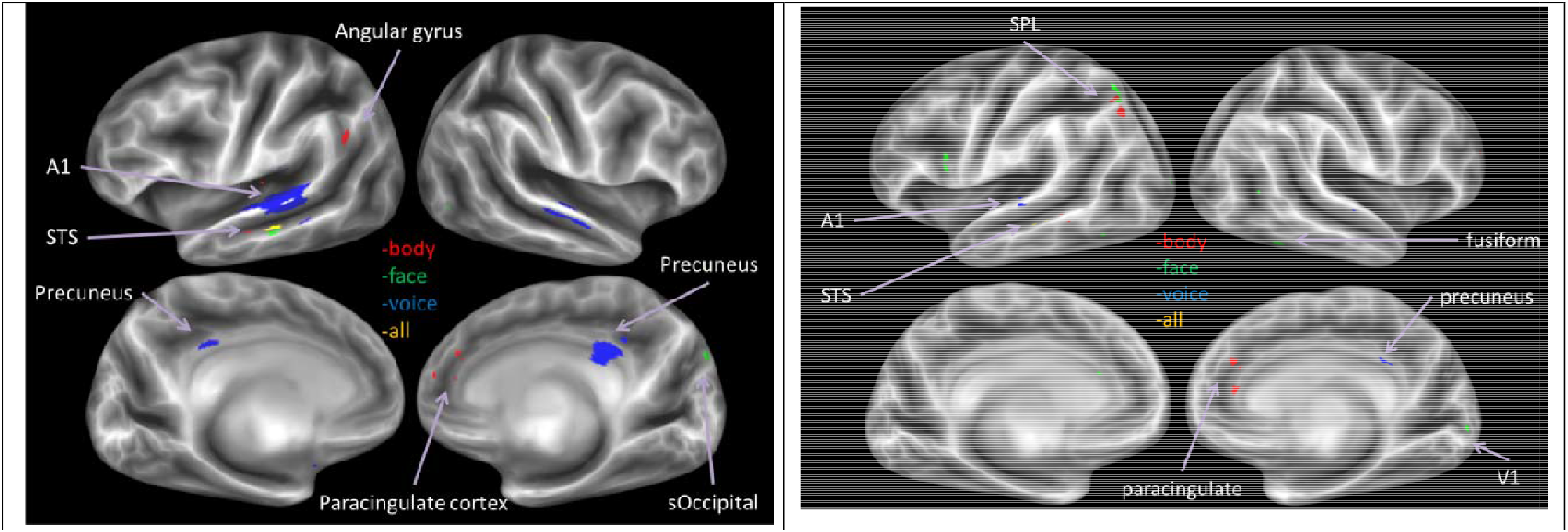
(Left) Decoder trained to classify emotion, for all stimuli and for each stimulus type separately each modality separately. (Right) Decoder trained to classify emotion excluding the neutral stimuli, for all stimuli and for each stimulus type separately each modality separately. The displayed results are label maps derived from the volume map thresholded at p<0.001 uncorrected.

**Supplementary table 1:**
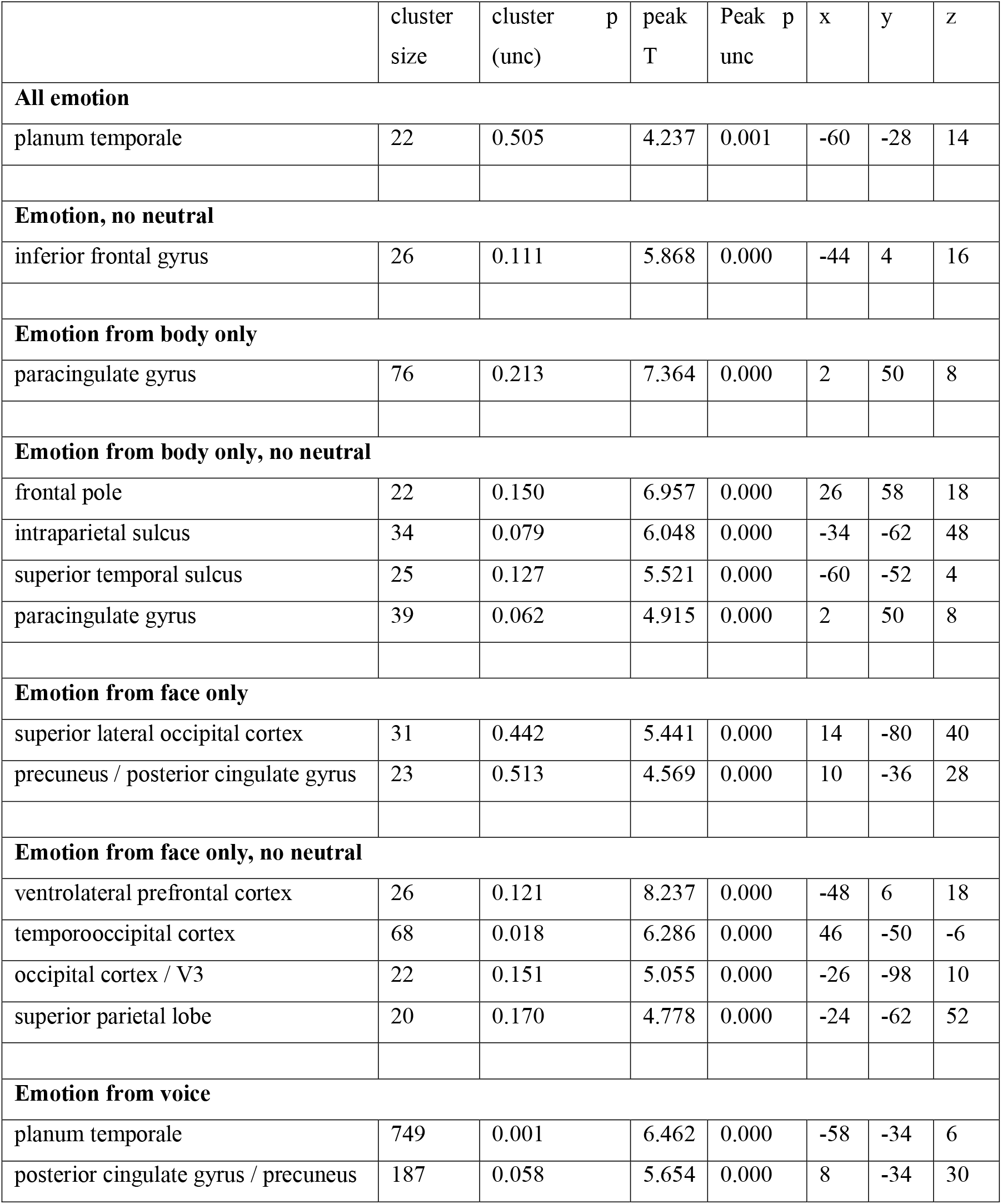
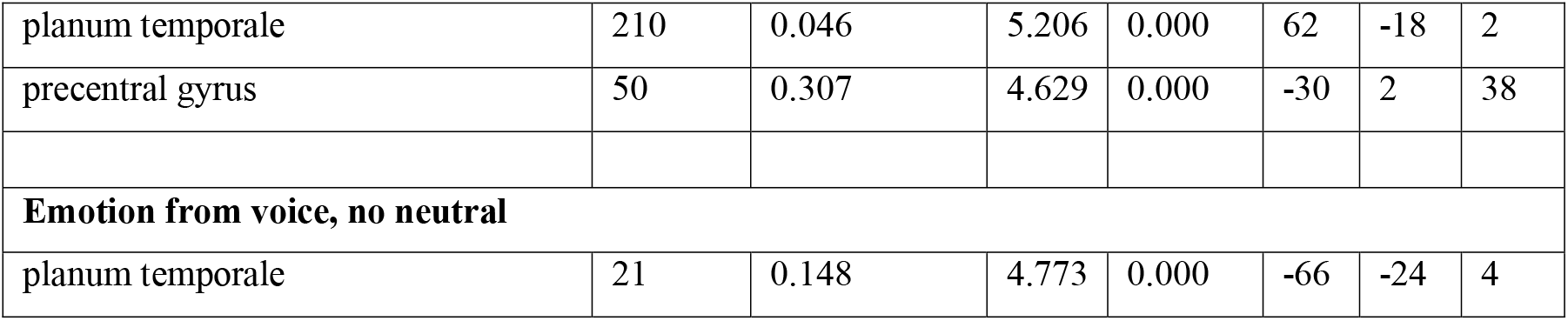

**Figure S5.**
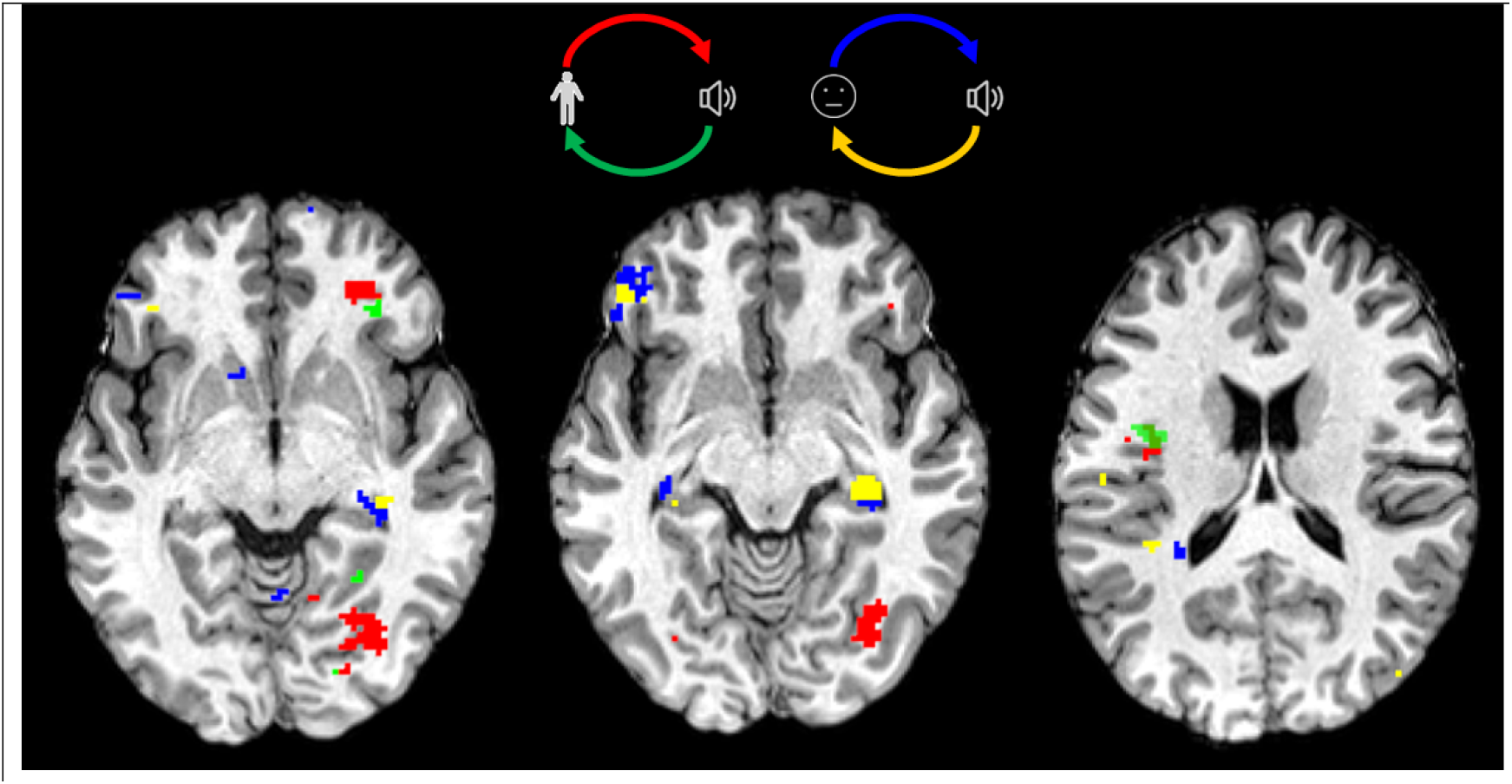
Crossmodal decoding of emotion. Red colors are for regions were emotion could be decoded from training on the body stimuli and tested on the voice stimuli from the body session. Green colors are for training on the voice stimuli from the body session and testing on the body stimuli. The same logic applies to stimuli from the face session (see top diagram). Results are at t>3.00 (p<0.01) uncorrected.

**Figure S6.**
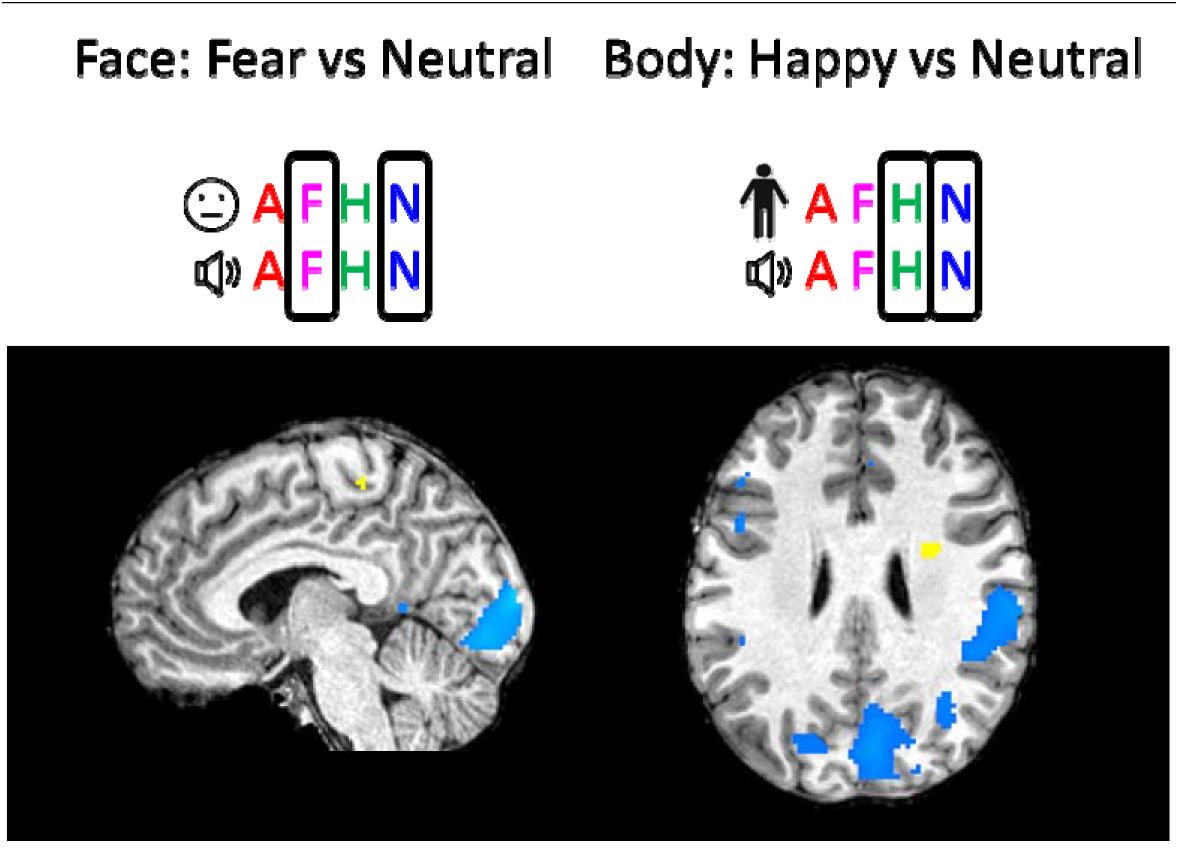
Comparison of decodability of modality and emotion. Results at p<0.001 uncorrected. Blue colors indicate where decoding accuracy for modality was higher than for emotion and vice versa for yellow colors. Diagrams above figures display what stimuli were used for decoding: the top left panel displays results for the face (face icon) and voice (sound icon) stimuli where the fear (F) and neutral (N) emotions were used and the angry (A) and happy (H) stimuli were left out.

